# Light-field deep learning enables high-throughput, scattering-mitigated calcium imaging

**DOI:** 10.1101/2025.03.17.643718

**Authors:** Carmel L. Howe, Kate L.Y. Zhao, Herman Verinaz-Jadan, Pingfan Song, Samuel J. Barnes, Pier Luigi Dragotti, Amanda J. Foust

## Abstract

Light field microscopy (LFM) enables volumetric, high throughput functional imaging. However, the computational burden and vulnerability to scattering limit LFM’s application to neuroscience. We present a light-field strategy for volumetric, scattering-mitigated neural circuit activity monitoring. A physics-based deep neural network, LNet, is trained with two-photon volumes and one-photon light fields. A processing pipeline uses LNet to extract calcium activity from light-field videos of jGCaMP8f-expressing neurons in acute cortical slices. The extracted time series have high signal-to-noise ratios and reduced optical crosstalk compared to conventional volume reconstruction methods. Imaging 100 volumes per second, we observe putative spikes fired at up to 10 Hz and the spatial intermingling of putative ensembles throughout 530 x 530 x 100-micron volumes. Compared to iterative algorithms, LNet workflows reduce light-field video processing times by 2- to 12-fold, advancing the goal of real-time, scattering-robust volumetric neural circuit imaging for closed-loop and adaptive experimental paradigms.

## Introduction

Calcium fluorescence imaging enables observation of neuronal network function over time scales of milliseconds to months, and elucidates structure-function relationships through the simultaneous measurement of position, morphology, and activity.^1^ Although calcium provides a low-pass filtered proxy for neuronal spiking well sampled at a few hertz, certain applications benefit from sampling faster. For example, functional connectivity maps can be inferred from the correlation of calcium signals imaged at 5 Hz, but we cannot infer the connection polarity, or whether connections are mono- or poly-synaptic.^2^ Hence next generation neural network monitoring must feature high speed, high sensitivity, and high throughput to capture the connectivity, dynamic states, and computations of living neural networks.

The development of fast, sensitive genetically-encoded calcium indicators (GECIs) and new optical imaging technologies have enabled the study of functional neuronal networks in a variety of organisms, including mammals. However, the goal of fast, high-throughput, and volumetric imaging poses a particular challenge in mammals as our brains scatter light strongly. This is because the modalities providing the best optical sectioning and scattering mitigation, such as confocal and two-photon (2P) laser scanning microscopy, have a limited fluorescence bandwidth due to scanning.^3^ The finite bandwidth limits the speed and number of cells that can be simultaneously monitored, and numerous strategies have been developed to boost or efficiently exploit the fluorescence generated.^4^ Rapid acquisition from selected regions of interest (ROIs) obtains in serial through optimized^5–7^ and rapid scanning and focusing,^8–15^ or parallelized through wavefront modulation.^16–23^ Scanned fluorescence excitation can be multiplexed and/or spatially extended into multifocal arrays,^24–36^ beads,^37–39^ lines,^40–46^ or planes.^47–54^ The added complexity of the above schemes generally limits their widespread use. Moreover, their bandwidth remains fundamentally limited to the extent of parallelization.

Light-field microscopy (LFM)^55^ is a fully scanless volumetric imaging modality which maximizes fluorescence bandwidth. A microlens array (MLA) is placed at the native focal plane of a widefield microscope, and the camera sensor is conjugated to the MLA back focal plane. The microlenses confer kilo-scale stereovision, enabling reconstruction of an entire volume from a single snapshot called the light field (LF). LFMs for calcium imaging are easily implemented with off-the-shelf MLAs, widefield fluorescence excitation, and complementary metal-oxide semiconductor (CMOS) cameras. Volumetric LFM calcium imaging was first demonstrated in transparent zebrafish larvae.^56–58^ In murine neocortex, LFM has achieved simultaneous recording of 1000s of neurons in a ø4 x 0.2 mm volume at up to 18 Hz^59^ and faster in smaller volumes.^60^ LFM’s maximal fluorescence bandwidth means that it will scale to image at faster rates and in more neurons thanks to faster, high pixel-count cameras and brighter, more sensitive calcium indicators.

Although LFM optical trains are simple and low-cost, to date, only a handful of neuroscience studies report their use. Two challenges hinder LFM’s widespread application to volumetric calcium imaging. The first is computational cost. Conventional model-based reconstruction of volumes from LFs rely on iterative 3D deconvolution.^61–63^ Recently data-driven deep learning approaches,^64–66^ and physics-inspired deep neural networks^67, 68^ have demonstrated fast, high-fidelity source localization and volume reconstruction in limited contexts. Certain approaches bypass volume reconstruction through phase space sparse coding,^69^ compressive sensing,^70^ or seeded-iterative demixing (SID).^60^ The second challenge is that most tissues, including the mammalian brain, scatter light strongly, while conventional LFM volume reconstruction assumes ballistic propagation. Scanned confocal,^71, 72^ speckle,^73^ selective volume,^74^ and other structured illumination LFM implementations^75–77^ mitigate scattered and out-of-focus fluorescence through spatial modulation of the excitation. Other strategies enhance signal extraction from scattering-corrupted LFs acquired with vanilla LFM optics through integration of scattering models,^78, 79^ compressive sens-ing,^70^ dictionary learning and convolutional sparse coding,^42, 76^ and iterative demixing.^60^ Structured illumination and post-processing strategies mitigate scattering effects and enable calcium extraction up to ∼400 microns in murine neocortex.^60^ Despite these advantages, LFM’s substantial computational burden precludes its application to activity-guided or closed-loop experimental paradigms.^80^

Deep-learning approaches stand to overcome LFM’s computational burden and enable real-time processing.^66, 81–83^ One key challenge for deep learning is ensuring reliability. A notable example for calcium imaging is HyLFM proposed by Wagner et al.,^65^ which integrates LFM with light-sheet microscopy, with the latter providing ground truth for training and continuous validation. We recently introduced two physics-based deep learning frameworks integrating the LFM wave-optics forward model and 2P scanned volumes to mitigate scattering effects while ensuring reliability of volume reconstruction^68^ and source localization.^67^ Deep learning integration of multimodal image acquisition can both mitigate scattering and accelerate calcium signal extraction.

Here we introduce a rapid LFM calcium activity extraction approach exploiting physics-based deep neural networks (DNNs) trained with one-photon light fields and a 2P scanned z-stack.^68^ We segment the active neurons from 100-Hz LF videos and compute each neuron’s calcium time series. LFs are imaged in mouse brain slices co-expressing the structural marker tdTomato and jGCaMP8f, the fastest available GECI, in neocortical layer 2/3 excitatory neurons. We compare the signal-to-noise ratio (SNR) and neuron-to-neuron optical crosstalk of calcium activity extracted by our DNNs to those extracted from volumes reconstructed by conventional Richardson-Lucy (RL) deconvolution and by seeded-iterative demixing (SID).^60^ Our pipeline extracts calcium activity from LF videos in FOVs for which the DNN has never viewed 2P volumes. Up to ∼60 active cells are detected simultaneously down to 100 microns below the slice surface. The 100 Hz LF capture and high SNR enable resolution of putative spikes fired up to 10 Hz. Compared to other methods, calcium activity segmented from the DNN-estimated volumes feature similar SNR, reduced crosstalk, and reduced computation time. We show that putative neuronal ensembles, i.e. groups with similar calcium activity, spatially intermingle throughout the LF-encoded volumes. Our strategy thus advances the goal of fast, high SNR, real-time volumetric calcium imaging necessary for speed-sensitive functional connectivity mapping, adaptive and closed-loop experimental paradigms. Importantly, the DNN generalizes to FOVs for which no 2P ground-truth are provided, making it possible to exploit 2P-like scattering mitigation without the 2P microscope.

## Results

### LNet enables scattering-mitigated calcium signal extraction from one-photon light-field videos

Light-field images acquired in mammalian brain tissues are blurred by scattering and fluorescence generated outside of the light-field encoded volume. We recently introduced a Learned Iterative Shrinkage-Thresholding Algorithm-based neural network (LNet) which estimates scattering-mitigated volumes from one-photon light fields.^68^ LNet first undergoes supervised training with a 2P scanned volume and a co-registered one-photon light field z-stack of layer 2/3 excitatory neurons transfected with the structural indicator tdTomato (Figure 1A). LNet then undergoes self-supervised training with light-field videos of the fast soma-localized calcium indicator ribo-jGCaMP8f. LNet integrates the LFM wave-optics forward model into the self-supervised training phase, which ensures reliable volume estimation. Indeed, good agreement is observed between the LNet output and the same volumes computed with purely model-based methods such as RL deconvolution (Figure 1B, see also^68^). Maximum intensity projections of one-iteration RL-deconvolved volumes feature low contrast and poorly confined neurons, similar to widefield^84^(Figure 2, Video S1). Contrast and confinement improve by increasing the RL deconvolution to eight iterations, but this also exacerbates the square-like artifacts around native focal plane. Strikingly, the LNet-estimated volumes feature 2P image characteristics including high contrast, well-confined neurons on a dark background. In addition, the LNet volume is free of the square-like artifacts characteristic of RL-deconvolved volumes.^63^

**Figure 1.**
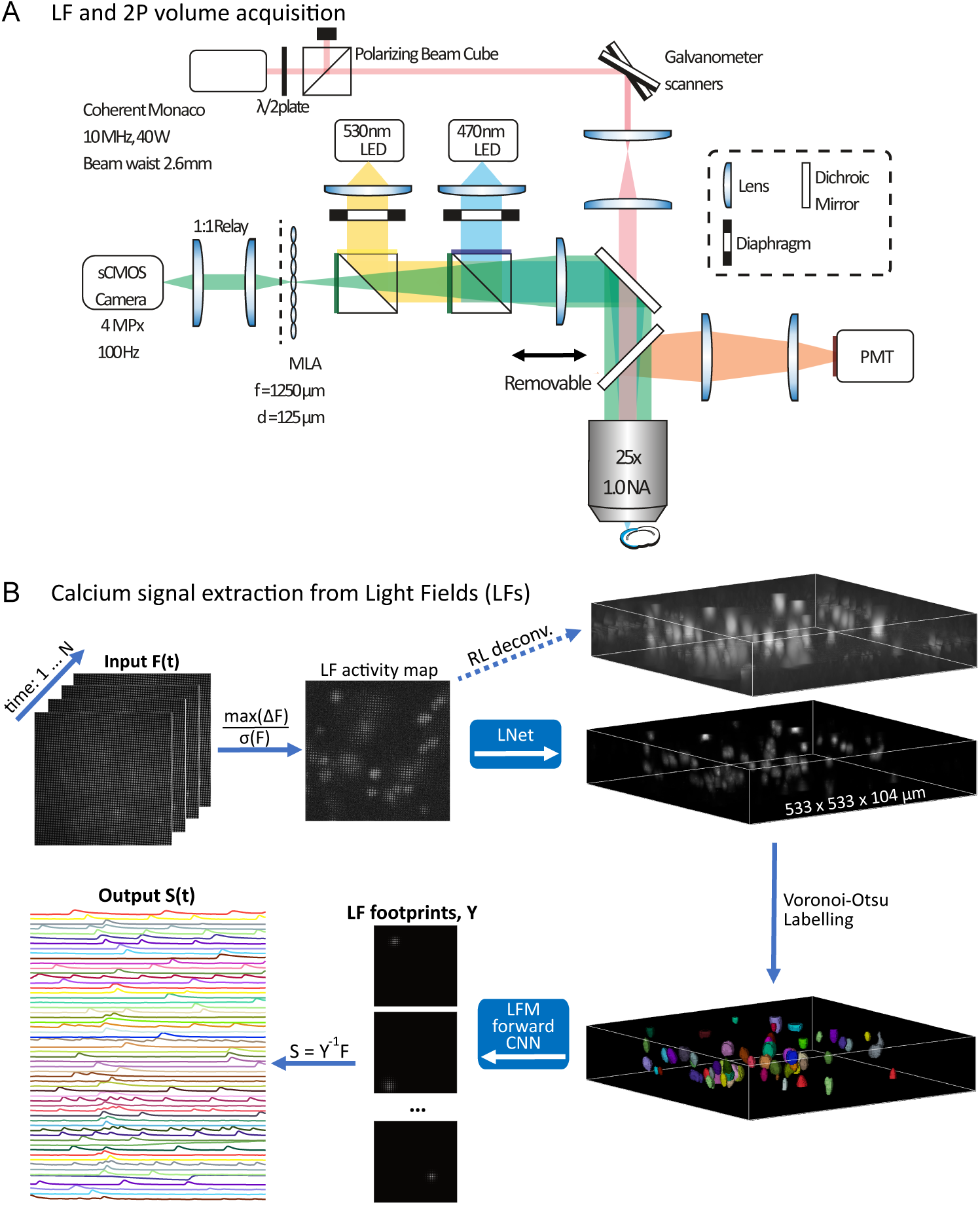
The LNet LFM and processing pipeline. (A) The dual-mode LFM combines a 2P laser-scanning path with a one-photon LFM. A removable dichroic mirror diverts fluorescence to the photomuliplier tube (PMT) for 2P collection. When the 2P dichroic is disengaged, fluorescence is imaged by the LFM with a microlens array (MLA). Distances not to scale. (B) Key elements of the deep-learning calcium activity extraction from one-photon light fields. Light field videos (F) undergo pixel-wise preprocessing (max(ΔF)/*σ*(F)) to generate a single light-field in which active neurons appear with high contrast. The light-field activity map forms the input to the LNet,^68^ which estimates a volume with 2P-like contrast and localization compared to the same volume computed via conventional 8-iteraton Richardson-Lucy (RL) 3D deconvolution. Neurons are segmented from the LNet-estimated volume with 3D Voronoi-Otsu labeling. Each of the ROIs separately forms an input to a CNN computing the LFM wave-optics forward model and hence each neuron’s “footprint” in the light field. The vectorized footprints form matrix Y, and the calcium time series (S) are found as the product of the pseudoinverse of Y and F, the light-field video.

**Figure 2.**
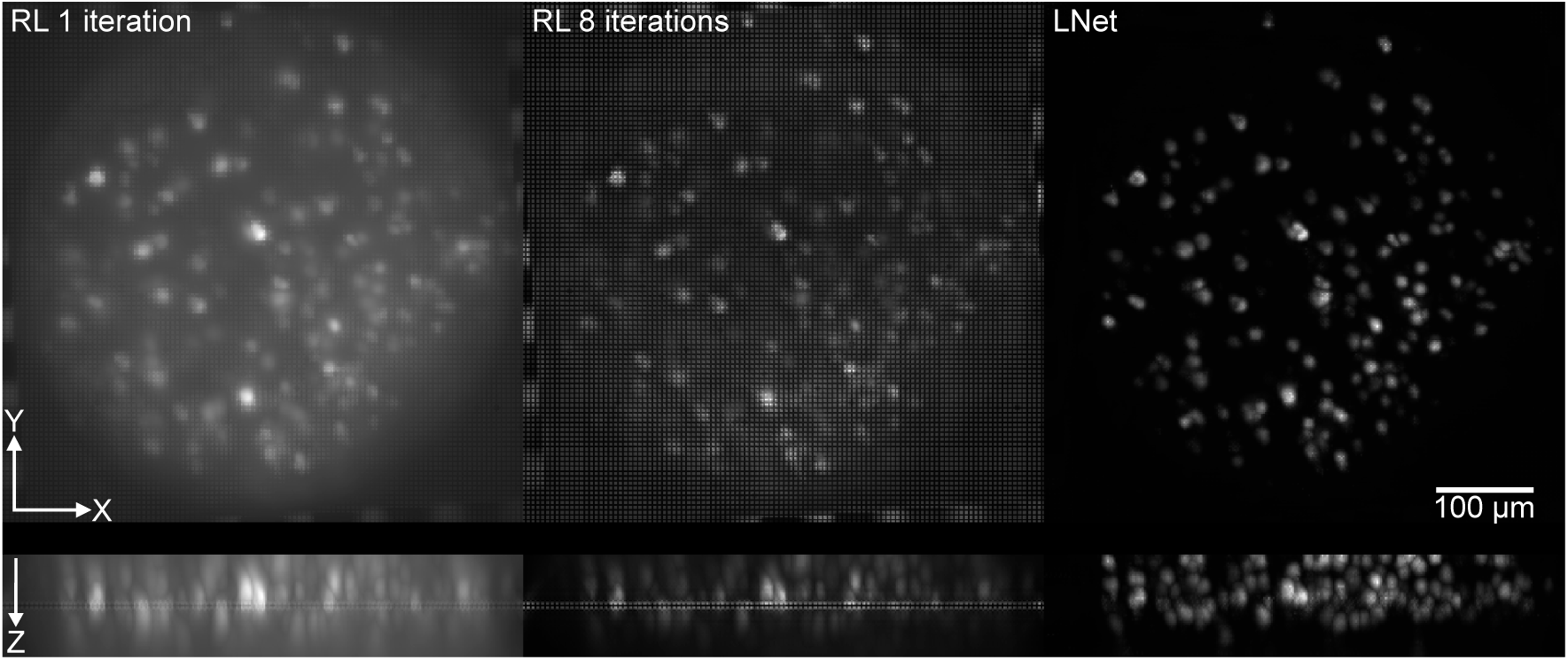
LNet estimates volumes from one-photon light fields with 2P-like background suppression and source confinement. The panels display x-y and x-z maximum intensity projections of volumes reconstructed via 1-iteration RL (left), 8-iteration RL (middle), and LNet (right) from a 100 Hz, five second light-field video of soma-localized jGCaMP8f fluorescence. See also Video S1.

We used LNet to systematically extract neuronal calcium activity from light-field videos via two methods. Both methods begin with a pixel-wise calculation of the maximum change in fluorescence (ΔF) divided by the standard deviation over time (*σ*(F)). The result is a single light field in which the active neurons appear with high contrast (Figure 1B). This light-field activity map forms the input to LNet. The LNet-estimated active neuron volume is segmented via Voronoi-Otsu labelling. In Method 1, called “LNet region of interest (ROI)”, LNet estimates volumes for each frame in the light-field video. The calcium time-series are then calculated as the mean intensity of voxels in each ROI of the activity map volume. In Method 2, called “LNet Matrix”, each ROI forms the input to a convolutional neural network (CNN) computing the LFM wave optics forward model, hence estimating each active neuron’s footprint in the light field. The footprints are vectorized and assembled into a matrix Y. The calcium time series are then computed as the product of the pseudoinverse of matrix Y and F, the light-field video. LNet LFM hence enables the rapid extraction of high signal-to-noise ratio (SNR) calcium transients from 100 Hz acquired light-field videos encoding 533 x 533 x 104 micron volumes (Figure 3). In this five second light-field video, ∼60 active cells were detected in a volume containing over 150 neurons.

**Figure 3.**
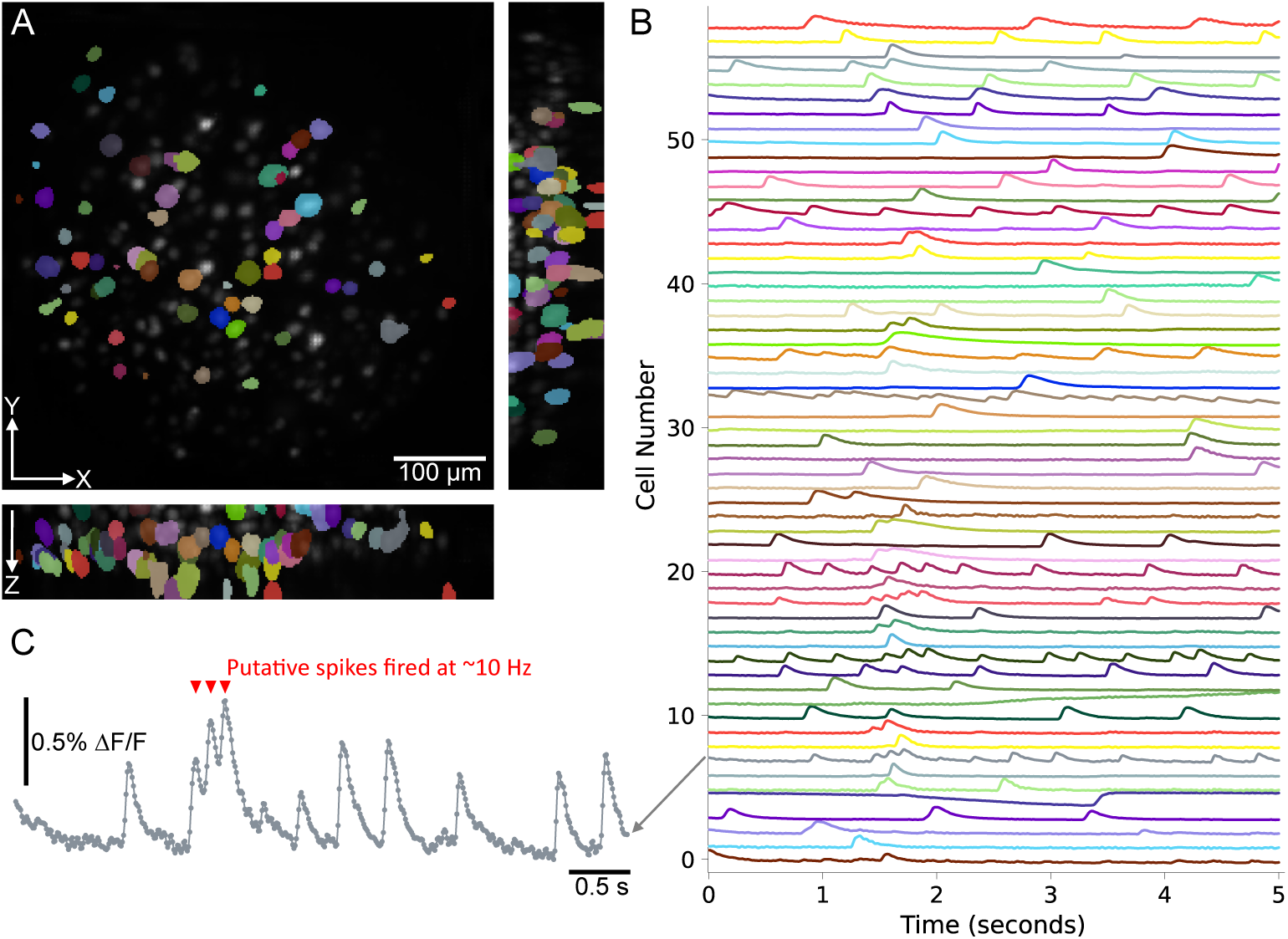
The LNet LFM enables the rapid extraction of high SNR calcium transients light-field videos. (A) Shows the maximum intensity projection of volumes estimated by LNet from a 100 Hz, five second light-field video. The overlaid ROIs are obtained with Veronoi-Otsu labeling of the volume estimated by LNet from the light-field activity map. The ROI label colors correspond to the same-colored time series obtained through matrix factorization displayed in (B). (C) shows an expanded view of one time series from (B). Red triangles indicate the time of putative spikes fired at a rate of ∼10Hz.

We benchmarked the SNR of time series extracted from 100 Hz jGCaMP8f light-field videos via LNet ROI, LNet Matrix, conventional 8-iteration RL deconvolution, and seeded iterative demixing (SID). SID is a state-of-the-art algorithm developed specifically to mitigate scattering effects in light-field calcium signal extraction.^60^ For RL, time series were extracted based on the LNet activity map ROIs, enabling paired comparison. As SID simultaneously localizes and demixes, the distribution of cells detected by LNet and SID differed, although with substantial overlap (Figure 4A). The time series extracted via LNet and RL were similar for each ROI (Figure 4B, Figure S1). The Pearson correlation coefficient of time series calculated for the same ROIs in the RL- and LNet-reconstructed volumes had a median of 0.99 and interquartile range (IQR) of 0.02 (n=107 cells). The high correlation with conventional model-based reconstructions reflects LNet’s reliability in this context. Both LNet and SID detected neurons throughout the 100-micron axial extent. LNet located 107 cells, and SID located 147 cells (time-series SNR *>* 6 dB, see Figure S2) across five 250-frame light-field videos acquired in five different FOVs (Figure 4C). SID detected more cells with low SNR compared to LNet across all depths (Figure 4D, E). The mean time-series SNR of LNet-localized cells was hence higher than SID-localized cells (Figure 4F). The RL-derived time series had the highest SNR (median = 13.2 dB, IQR = 2.4 dB), followed by LNet Matrix (median = 13.0 dB, IQR = 2.6 dB), LNet ROI (median = 12.7 dB, IQR = 2.8 dB), and SID (median = 10.9 dB, IQR = 5.2 dB) across 5 FOVs obtained from 4 cortical slices in 3 mice (Figure 4F). We identified matched time series for SID and LNet as cells with centers separated by less than 20 microns and time-series correlation coefficients *>* 0.5 (n = 75 neurons from five FOVs). The matched comparision showed RL-derived time series SNRs significantly higher than those obtained with the other methods (as median, IQR in dB; RL: 13.4, 2.8; LNet Matrix: 13.0, 2.8; LNet ROI: 12.8, 3.0; SID: 12.6, 4.0; Figure 4G). In summary, LNet enables extraction of high SNR calcium time series.

**Figure 4.**
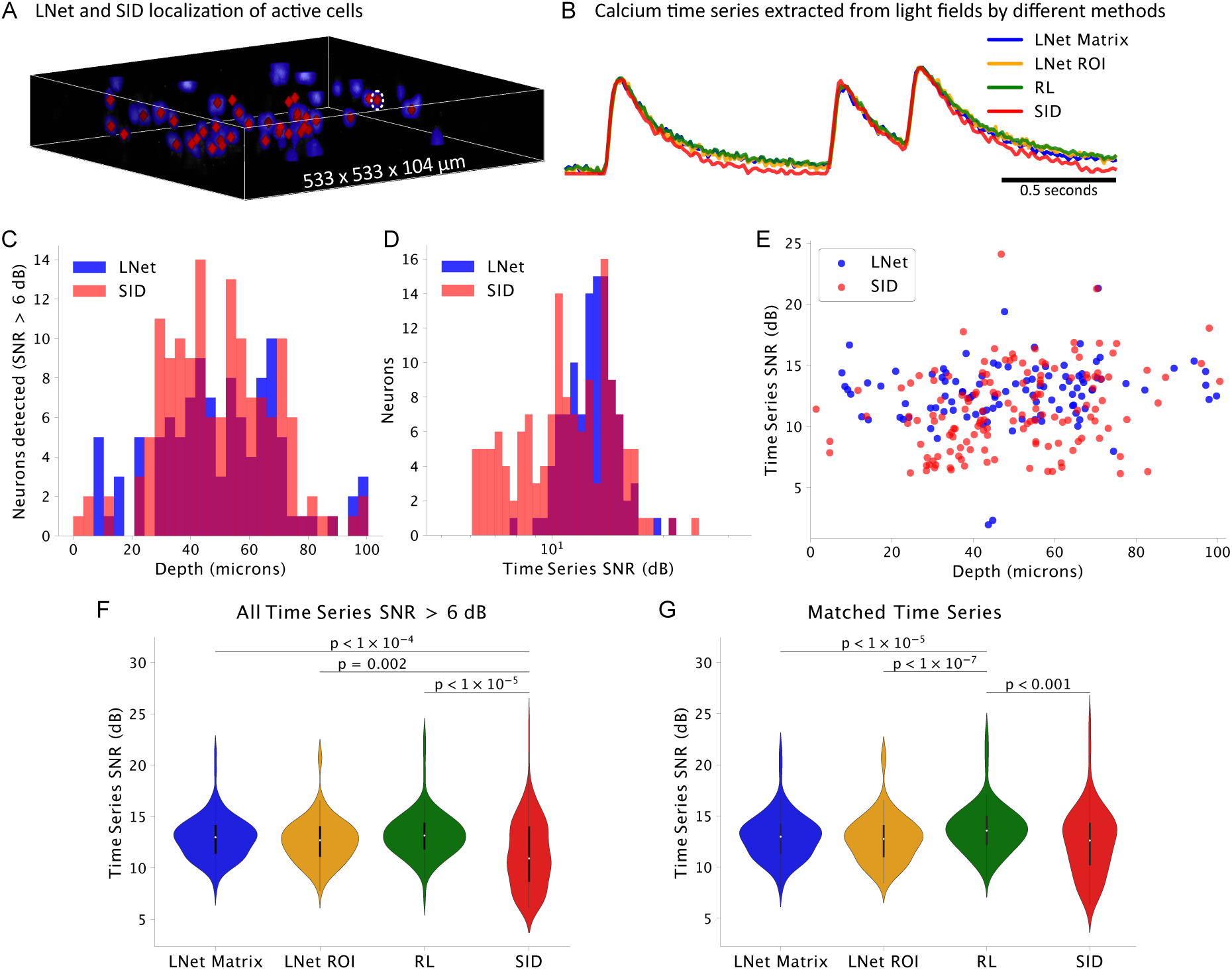
The LNet LFM detects high signal-to-noise ratio calcium signals. The SNR of LNet-extracted calcium signals benchmarked to conventional 8-iteration Richarson-Lucy (RL) deconvolution and seeded-iterative demixing (SID^60^). (A) shows the LNet estimated activity map for a single ROI overlaid with LNet-segmented ROIs (blue) and neuron centroids localized by SID (red diamonds). (B) displays the normalized calcium time-series for the neuron outlined with a dashed white line in (A), extracted via four methods: (1) LNet Matrix, (2) LNet ROI, (3) RL deconvolution, and (4) SID. (C) shows the number of neuronal calcium time series detected with SNR *>* 6 dB via LNet segmentation (blue) and SID (red) as a function of depth. (D) displays the histogram of SNRs for the same neurons (all depths) and (E) plots the SNR as a function of depth. (F) compares the SNR for all neurons with calcium time-series SNR *>* 6 dB across methods (independent samples t-test, p-values Bonferroni-corrected for multiple comparisons). (G) Displays SNR for “matched” time series in which neuronal centers are *<*20 microns apart and the Pearson correlation coefficient is *>* 0.5 (paired samples t-test, p-values Bonferroni-corrected for multiple comparisons).

### LNet reduces calcium signal cross-talk in neighboring neurons

Light scattering and insufficient optical sectioning cause crosstalk between calcium signals arising from adjacent neurons. Figure 5A shows two axially adjacent cells, a worst-case configuration for optical crosstalk. Since point spread functions are extended in the axial dimension, signals arising from laterally overlapping and axially adjacent sources, such as the neurons indicated by the red and blue ROIs, are the most difficult to unmix. Indeed, the time series extracted from these neurons via conventional 8-iteration RL deconvolution of each volume in the light-field video show substantial crosstalk (Pearson correlation coefficient = 0.60). Time-series from the same ROIs in the LNet-estimated volumes show substantially less crosstalk, with a correlation coefficient of 0.41, two-thirds that of the RL-reconstructed volume series.

**Figure 5.**
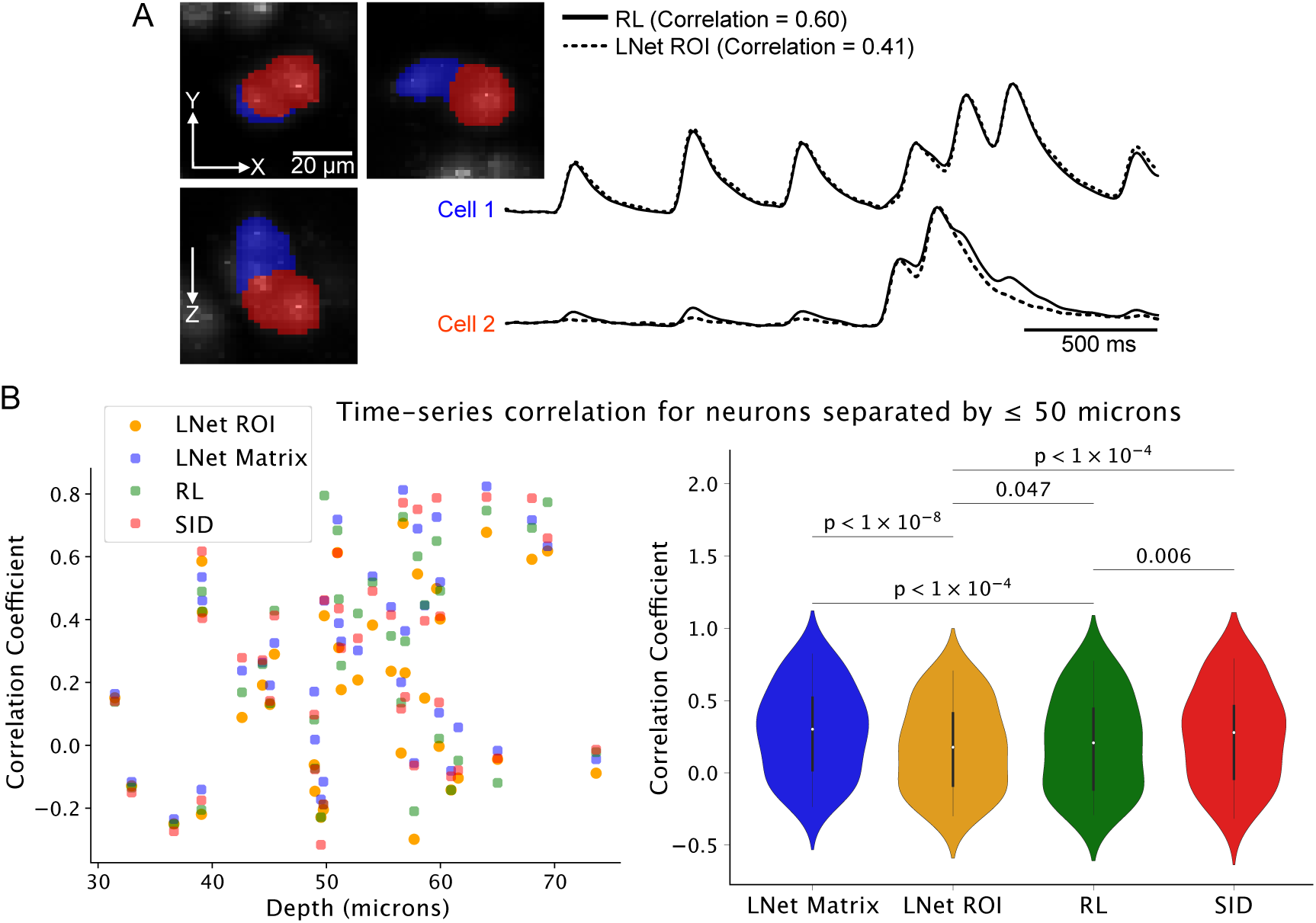
Calcium time-series extraction from LNet-reconstructed volumes minimizes optical crosstalk. (A) Two cells segmented in the worst-case scenario for optical crosstalk: overlapping laterally and touching axially. The time-series extracted from these ROIs in RL-deconvolved volumes (solid black traces) are highly correlated (coefficient = 0.60) reflecting high crosstalk. Extracting the time-series from the same ROIs in the LNet-estimated volume series (dashed black lines) decreases this correlation (coefficient = 0.41). (B) Pearson correlation coefficients of calcium time series extracted via deep learning (“LNet ROI”, “LNet Matrix”), 8-iteration RL deconvolution, and SID. Time series were matched across methods, and the correlation between neighboring (centroids separated by ≤ 50 microns) neuron time-series is plotted as a function of the mean pair centroid depth. (C) Comparison of neighboring cell correlation coefficients for time series extracted via four methods (n=39 neighboring neuron pairs across five FOVs, paired samples t-test, p-values Bonferroni-corrected for multiple comparisons).

We benchmarked LNet’s neuron-to-neuron crosstalk against RL deconvolution and SID.^60^ We calculated the correlaton coefficient of calcium signals from of cells separated by a euclidean distance of ≤ 50 microns. Figure 5B shows the correlation of matched time series as a function of the mean depth of the pair of neurons (n = 39 neighboring neuron pairs from five FOVs). Time series from nearby neurons extracted with the “LNet ROI” method had lower correlation (median = 0.18, IQR = 0.50) than those extracted with the “LNet Matrix” method (median = 0.30, IQR = 0.50), RL (median = 0.21, IQR = 0.56), or SID (median = 0.28, IQR = 0.50; Figure 5B). Although such correlations arise from a combination of optical crosstalk and true connectivity, the difference in correlation across these for methods in matched adjacent neurons reflects differing levels of optical crosstalk. The “LNet ROI”-extracted time series featured the lowest crosstalk across mean pair depths up to 74 microns.

### LNet LFM’s speed and sensitivity volumetrically resolves putative spikes fired at 10 Hz

We exploited LNet LFM’s bandwidth to track neuronal spiking dynamics reported by the fastest available GECI, jGCaMP8f,^85^ in volumes at 100 Hz. In our dataset, this enabled high SNR tracking of putative spikes fired at rates up to ∼10 Hz (Figure 3C). Typically, volumetric and multiplane calcium imaging is performed at only a few Hz. Figure 6 shows the impact of downsampling jGCaMP8f time series to 50 Hz, 20 Hz, 10 Hz, and 5 Hz. Here we perform the downsampling by averaging, hence boosting the SNR for lower rates by the square root of the downsampling factor. We detected putative spikes as peaks in the deconvolved calcium traces.^86^ Even though averaging boosts the 5 Hz and 10 Hz time series SNR by 3-to 4-fold, putative spikes are missed and the timing of detected spikes distorted.

**Figure 6.**
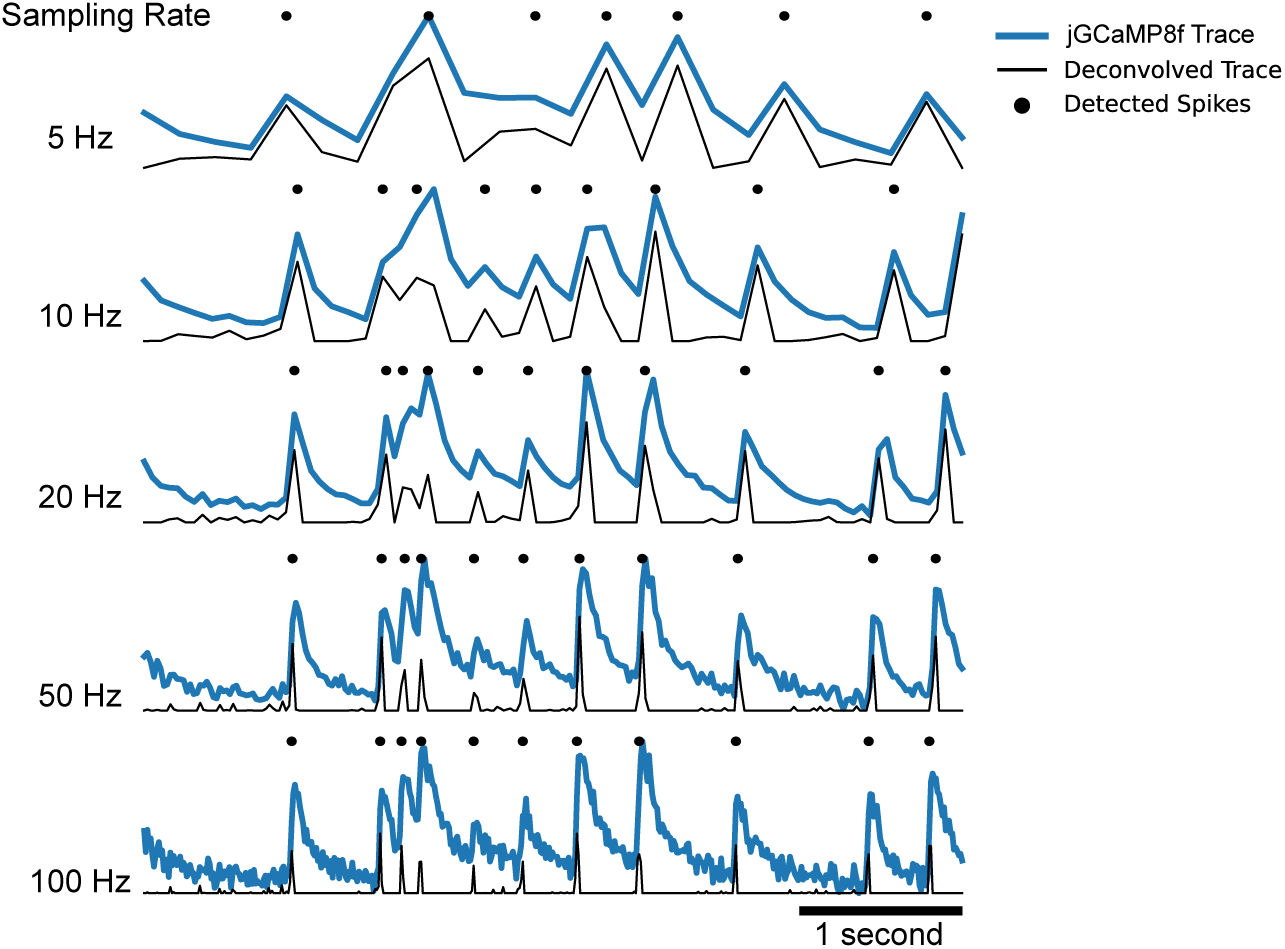
Deconvolution of neuronal calcium signals obtained with LNet LFM at 100 volumes/second enabling detection of putative spikes fired at 10-Hz rates. Example jGCaMP8f calcium time series (blue traces) are deconvolved using the OASIS package (black traces).^86^ Peaks in the deconvolved traces mark the timing of putative spikes (black dots). Downsampling by averaging adjacent time points boosts SNR but results in missed spikes and distorted spike timing.

### Neuron ensembles spatially intermingle in LFM-encoded volumes

The high speed and SNR of LNet LFM can be exploited to map neuronal ensembles in 3D. We performed agglomerative hierarchical clustering on the deconvolved 100-Hz calcium time series extracted with LNet from each of five FOVs. Figure 7 shows the calcium time series (A) and Pearson correlation coefficients (B) for the clustered deconvolved calcium signals extracted from one of the FOVs. The euclidian distance between cells within a cluster (median = 191 microns, IQR = 158 microns, n = 579 distances) and between clusters (median = 193, IQR = 137 microns, n = 2079 distances) was not statistically different (Figure 7C). Indeed visual inspection of the segmented cell clusters in the example FOV shows that cells with correlated activity spatially intersperse among cells from other groups throughout the volume (Figure 7D).

**Figure 7.**
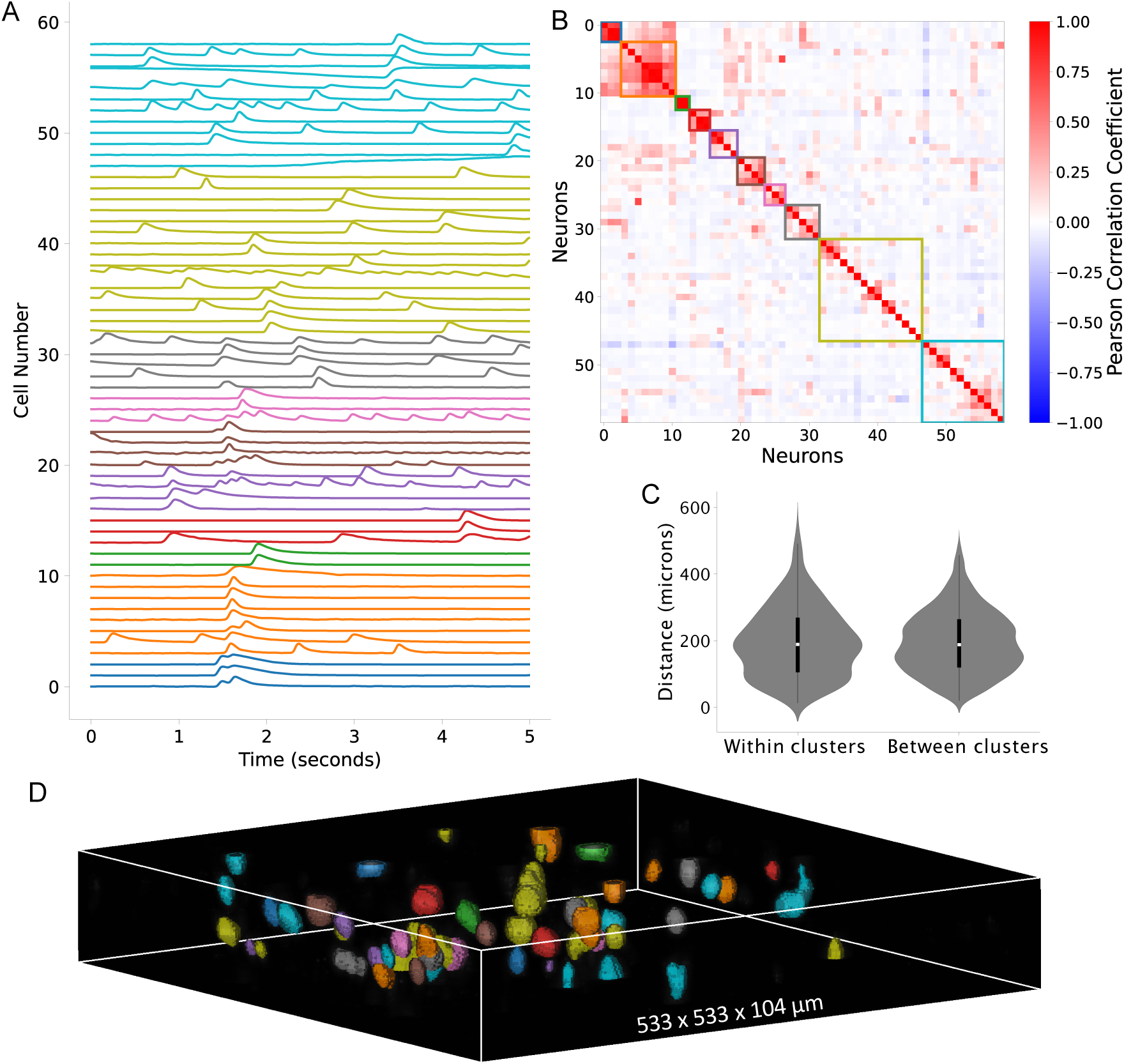
Neurons with correlated calcium activity intermingle in LNet LFM-encoded volumes. (A) Calcium time series and correlation matrix of the deconvolved traces (see Figure 6) ordered and grouped with agglomerative hierarchical clustering. The color-coded time series and boxes on the correlation matrix indicate the clusters. (C) The euclidian distance between cells within and between clusters from five FOVs was not statistically different (p *>* 0.5 for Mann-Whitney U and independent samples t-tests). (D) visual inspection of color-coded ROIs in the volume shows spatial intermingling of neurons from different clusters.

### LNet reduces single-workstation light-field video processing from hours to minutes

The computational load and complexity of volume reconstruction from light fields has previously hindered LFM’s application to neurophysiology and biological imaging in general. Nöbauer et al. previously demonstrated that SID reduced computational resource requirements by three orders of magnitude compared with conventional LFM reconstruction (e.g. iterative 3D deconvolution such as RL), bringing LFM calcium videos processing down to the capacity of a single workstation.

We compared the computation time required for LNet-based calcium time-series extraction with that of SID on a single workstation equiped with a NVIDIA GeForce RTX 2080 Ti graphical processing unit (GPU). Calcium time-series extraction with SID took ∼12 hours per 250-frame light-field video. Initial training of LNet took ∼20 hours (400 epochs at ∼3 minutes/epoch). Following this one-time training, LNet was applied to jGCaMP8f videos in previously unseen FOVs with brief fine-tuning taking ∼30 minutes (10 epochs at ∼3 minutes/epoch) with the LF activity maps for the “LNet Matrix” workflow. For the “LNet ROI” method, LNet was fine-tuned simultaneously on every 6th frame of the five light-field videos for 100 epochs. This fine tuning took 300 minutes or five hours (100 epochs at ∼3 minutes/epoch). Subsequent light-field preprocessing, volume reconstruction, segmentation, and time-series extraction, either via footprint estimation and matrix factorization (“LNet Matrix”) or by averaging intensities from the LNet-estimated volume series (“LNet ROI”) took only a few minutes. Hence, although the initial computationally greedy LNet training takes several hours, subsequent calcium signal extraction from unseen FOVs using the “LNet Matrix” workflow takes under 40 minutes, including fine tuning, on a single workstation. “LNet ROI” required more time for fine tuning but reduced optical crosstalk compared to all methods tested.

## Discussion

In this article we have introduced physics-based deeping learning strategies for high sensitivity, scattering-mitigated volumetric calcium signal extraction from light field videos. The videos were acquired at 100 Hz, enabling high-speed volumetric capture of neural dynamics reported by the fast GECI jGCaMP8f. The LNet, trained on a single 2P volume stack and fewer than 300 one-photon light fields, conferred 2P-like contrast and source confinement to volumes estimated from LFs encoding previously unseen FOVs. The LNet LFM enabled extraction of calcium time series within a ∼530 x 530 x 100 micron volume in mouse neocortical brain slices. Ensembles of neurons with similar calcium time series spatially intermingled throughout the volumes.

One primary challenge with deep learning is ensuring reliability. DNNs can hallucinate features from noise. The integration of the wave-optics forward model into LNet’s architecture protects against unrealistic volume estimation. Indeed, time series extracted from the the same ROIs in the LNet- and RL-reconstructed volumes were highly correlated (median coefficient 0.99) indicating high reliability.

LNet LFM resolved putative spikes fired at rates up to 10 Hz. Its high volume rate and SNR, combined with jGCaMP8f’s fast decay,^85^ can improve tracking of complex spike events and of interneurons which produce smaller calcium transients, higher spike rates, and less bursting than excitatory neurons.^87^ Moreover, the 100 Hz volume rate captured 3-4 points on the fast rising phase of each calcium transient (Figure 3C), adequate to infer connection polarity with 10 millisecond precision. The volume rate was limited by the camera’s maximum full field frame rate. Faster, more sensitive and low-noise sCMOS cameras are already commercially available, which would enable even faster volume rates with an approximately 1.5 dB SNR loss per speed doubling all other factors being equal.

Compared to scattering mitigation strategies based on scanned or structured illumination,^3^ scanless LNet LFM stands out because of its maximal fluorescence bandwidth, speed (100 volumes/s) and sensitivity (median SNR ∼13 dB). Compared to LNet, the SID algorithm detected more time series with SNR *<*10 dB than LNet. SNR was similar for time series detected by both strategies. While 8-iteration RL achieved the highest SNR of all methods, it had worse optical crosstalk than the LNet ROI pipeline. We used ROIs segmented from the LNet-estimated activity volume to extract the time series from the RL-reconstructed volumes. Hence, this comparison likely quantifies the best case for RL crosstalk. In practice, high background and poor source confinement make it difficult to segment the RL-reconstructed volumes on their own.

One striking feature of LNet is the ability to confer 2P-like contrast and source confinement to FOVs for which it only “sees” one-photon light fields. Indeed, time series extracted from the same neighboring neuron pairs from the LNet-reconstructed volumes were less correlated than those extracted from RL-reconstructed volumes, implying reduced optical crosstalk. The LNet ROI method also reduced crosstalk compared to SID, which discriminates calcium activity for neurons separated by distances down to 20 microns according to ground truth 2P imaging.^60^ Indeed, minimizing optical crosstalk between nearby neurons is critical for the accurate characterization of fine spatial scale functional motifs such as microclusters.^88^

Following the one-off computationally greedy LNet training, LNet can reliably extract calcium time series from previously unseen FOVs following brief fine tuning. The entire “LNet Matrix” pipeline, including fine-tuning, takes under 40 minutes on a single work station, an 18-fold reduction compared to SID processing of the same videos. Fine-tuning for the “LNet ROI” pipeline took ∼5 hours and achieved the lowest optical crosstalk. There is scope to reduce the time needed for fine tuning by, e.g., keeping only a few layers trainable or adjusting the loss function. Further adaptation of LNet to GPU and field-programmable gated array (FPGA) architectures^89^ may soon enable real-time signal extraction for adaptive and closed-loop neuroscience experiments.

In this study we imaged the most superficial 100 microns of submerged neocortical brain slices, which contained the well-perfused active cells. Future studies will assess LNet’s performance at greater depths in-vivo. In-vivo imaging will require correction of motion artifacts not encountered here in-vitro. For motion correction, widefield LFMs have a significant advantage over scanned or structured illumination strategies where motion can cause complete signal loss as sources shift out of the excitation pattern. Widefield LFMs excite and encode fluorescence simultaneously throughout the volume which can be aligned post hoc.

The initial training of LNet was performed with a single 2P z-stack. Application to other FOVs required no additional 2P data. It is conceivable that LNet could also process light fields acquired by other microscopes with analogous LFM optics without the costly 2P laser, scanning and collection optics. Our particular LFM’s resolution and ∼530 x 530 x 100 micron FOV were determined by our microscope objective and MLA, but in principle bespoke versions of LNet can be trained for a variety of LFMs, including mesoscale^59^ and head-mounted miniscope^90^ configurations, given a small set of images acquired in both 2P and one-photon LFM modes. The advantage is that, following the appropriate one-off training, end users would only need simple LFM optics and a single work station. This would enable access to high throughput, scattering-mitigated, volumetric imaging capability with simple hardware similar to widely used widefield imaging systems like Inscopix. A critical next step to realizing this vision is characterizing LNet’s generalizability to different LFM optics, including system-to-system alignment variation, and to samples expressing different indicators at different sparsities and depths.

## Methods

### The dual mode one-photon light-field and 2P laser-scanning microscope

The home-build upright microscope collected 2P scanned images and one-photon epifluorescence light fields in the same FOVs (Figure 1A).

For 2P imaging of tdTomato, a large removable dichroic mirror (50 × 70 × 2 mm, T750lpxrxt-UF2, Chroma) engaged directly above the objective (25x, NA 1.0, XLPLN25XSVMP, Olympus) back aperture, which was conjugated and demagnified onto the active area of a photomultiplier tube module (PMT, Hamamatsu H10722-20-10MHz). The tdTomato fluorescence was shortpass filtered with a 750-nm cutoff (Semrock FF01-750/SP-25). The PMT was powered by a programmable

DC power supply (Radiospares, RSPD3303C) controlled by a custom graphical user interface GUI written in Python 3. A Mini-circuits BLP-1.9+ (2.5 MHz cutoff) low-pass filtered the PMT module output voltage. The conditioned PMT signal was digitized by a National Instruments PCI-6110 and assembled into images and z-stacks using ScanImage version 3.8 software. ScanImage also controlled the scan engine through two PCI-6110 analog outputs to high-power servo boards (671315K-1HP, Cambridge Technologies) driving three-millimeter galvonometric mirrors (6M2003S-S, Cambridge Technologies) which laterally scanned the beam of a Coherent Monaco amplified ultrafast fibre laser (Coherence Monaco 1035-40-40, center wavelength 1035 nm, pulse width 286 fs, frequency 10 MHz, beam waist 1/e^2^ 2.6 mm, 5-10 mW power under the objective). A 30-millimeter focal length scan lens and a 300-millimeter focal length tube lens conjugated and expanded the beam at the galvos onto the objective back aperture. A dichroic mirror (DI03-R785-T3 25 × 36 × 3 mm, Semrock) between the tube lens and the 2P collection dichroic coupled the beam into the optical train shared with the one-photon LFM path.

For the LFM, one-photon fluorescence was excited in widefield with light-emitting diodes (LEDs) conjugated to the objective back aperture with 35-millimeter focal length plano-convex lenses and a 180-millimeter tube lens (TTL180-A, Thorlabs). TdTomato fluorescence was excited with a 530 nm LED, and jGCaMP8f fluorescence was excited by a 470 nm LED, both powered by an OptoLED current driver (P1110/002/000, Cairn Research). Semrock FITC-Ex01-Clin (excitation), Semrock FF495-DI03 (dichroic), and Semrock FF01-550/88 (emission) formed the jGCaMP8f filter set. Semrock FF01-520/35 (excitation), Semrock FF552-Di02 (dichroic), and Chroma ET605/52m (emission) formed the tdTomato filter set. The objective and the tube lens imaged the native focal plane onto an off-the-shelf microlens array (MLA, 125 µm pitch, f/10, RPC Photonics). A 1:1 macro lens (Nikon 60 mm f2.8 D AF Micro Nikkor Lens) relayed the MLA back focal plane onto the scientific CMOS camera (ORCA Flash 4 V2 with Camera Link, 2048 × 2048 pixels, 6.5 µm pixel size, Hamamatsu).

The LFM native lateral resolution is approximated by the microlens pitch (125 µm) divided by the objective magnification (25x), or 5 µm. The axial resolution of a LFM is determined by the number of resolvable diffraction-limited spots behind each microlens.^55^ Using the Sparrow criterion and assuming an average emission wavelength of 530 nm, the spot size in the camera plane is 6.2 µm, slightly smaller than the 6.5-µm sCMOS camera pixels. Hence, given the 125-µm MLA, we are able to resolve ∼19 distinct spots under each microlens. This corresponds to a synthetically refocusable depth-of-field of 7.5 µm over a 131-µm axial range. The deconvolved resolution generally exceeds the above in a depth dependent fashion outside of the native focal plane where angular information is redundant and square-like artifacts dominate.^63^

Light-field videos of jGCaMP8f-expressing neurons were imaged at 100 frames/second using Micromanager 2.0-gamma.^91^ A stepper motor (SliceScope, Scientifica) controlled by ScanImage and Micromanager scanned the objective along the optic axis to acquire 2P and light-field z-stacks of tdTomato-expressing neurons.

### LNet structure and training

We recently introduced LNet as a means to exploit the scattering mitigation of 2P laser scanning microscopy to estimate low-background, well-sectioned volumes from blurry one-photon light-fields. Full details of LNet architecture and training can be found in reference^68^. Briefly, a filter bank representation of the LFM wave-optics forward model is expressed as a CNN. We formulate the inverse problem, inferring volumes from light fields, as an optimization promoting sparsity in the reconstruction, since the compact jGCaMP8f somata occupy a small fraction of the total reconstructed volume. The optimization is solved using the Iterative Shrinkage-Thresholding Algorithm whose iterations are unfolded into the layers of a CNN, the LISTA network or “LNet”. The parameters of LNet are trained in two phases. The first phase consists of supervised training with a light-field z-stack (28 images with a 2-micron z-step) and a 2P z-stack (80 planes with a 2-micron z-step) of the same tdTomato expressing neurons in a cortical brain slice. Each light field in the z-stack is paired with a 2P substack consisting of 53 depths, 2-micron step, vertically centered on the LFM native focal plane. This results in 28 light-field/2P-volume pairs which form the labeled data set for supervised training. The second training phase is self supervised, since there are no ground truth 2P volumes for the jGCaMP8f light field videos. Here the LFM forward model CNN computes light fields from volumes estimated by LNet, and the loss between the computed and input light fields is minimized. The light-field loss is regularized by an adversarial loss consisting of a critic trained to distinguish between real and false volumes based on the 2P z-stack. The self-supervised training was conducted with three 80-frame jGCaMP8f videos acquired at 50 frames/second from three different FOVs.

In total, LNet was trained with very little data: 108 structural images (a 28-image light-field z-stack and an 80-image 2P z-stack from a single FOV) and 240 jGCaMP8f light-field video frames (three 80-frame, 50 Hz videos). Impressively, LNet generalizes to previously unseen FOVs, for which there is no 2P ground truth, including the full dataset presented here, with only brief fine tuning. For reconstructing the five light-field activity volumes, fine tuning was run for 10 epochs using five light-field activity maps calculated from the five light-field videos. For the LNet ROI pipeline, we fine tuned LNet for 100 epochs on every 6th frame of the five light-field videos simultaneously. Importantly, fine tuning retains the original training data while incorporating only the new jGCaMP8f light-field images, without requiring additional modalities or labeled datasets.

### jGCaMP8f time-series extraction from light-field videos

Our time-series extraction approach starts with preprocessing of jGCaMP8f light-field videos to generate a single light-field activity map in which active neurons appear with high contrast (Figure 1B). First, a size three uniform filter is applied to each pixel across time. Max(ΔF) is calculated in each pixel and divided by the standard deviation of the unfiltered time series, *σ*(F). The Max(Δ F)/*σ*(F) array is then multiplied by a factor of 1024 and the median pixel value calculated. The difference of the median of the scaled array and 16 is subtracted from each pixel. Then all values are truncated to a maximum value of 150 (such that higher values are set to 150) and a minimum value of 9, corresponding to the camera dark value on an 8-bit scale. These pre-processing steps make active neurons appear with high contrast in the activity map while keeping the minimum, maximum, and median values similar to those of the jGCaMP8f LNet training videos. Light-fields were rectified and formatted in Matlab with a custom script.

The light-field activity map forms the input to LNet which estimates the active neuron volume of size 231 x 231 x 53 voxels covering a 533.3 × 533.3 × 104-micron volume. The activity map volumes are segmented with the Voronoi-Otsu labeling method (py-clEsperanto prototype, *σ_spot_* = 3 and *σ_outline_* = 1). From this the “LNet ROI” and “LNet Matrix” pipelines diverge. For “LNet ROI”, LNet is fine-tuned on every sixth frame of the light-field video and then reconstructs the entire volume series. Time series are calculated as the mean intensity of pixels across time for each ROI in the segmented activity volume. For “LNet Matrix” each segmented ROI is put into an 8-bit volume containing zeros everywhere except for voxels within the ROI which are set to a value of 150. Each ROI volume is then fed to the LFM forward model CNN which computes each neuron’s contribution, or “footprint”, in the light field. The neuronal light-field “footprints” are flattened and assembled into a matrix Y. The calcium time-series (S) are then obtained through spatiotemporal factorization:

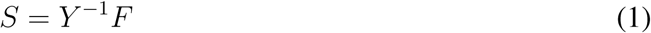

where F is the light-field video and *Y*^−1^ is the pseudoinverse of the rectangular light-field footprint matrix. The LNet processing pipelines are implemented in Python 3.

### Implementation of benchmark methods

We benchmarked the performance of the “LNet Matrix” and “LNet ROI” methods against that of conventional model-based 3D RL deconvolution and SID.^60^

For RL, we reconstruct each frame individually to obtain a volume time series. We used 8 iterations with the backprojection volume from the transpose operation as the initialization. Time-series were calculated as the average intensity of each ROI across the volume time series, using the ROIs segmented from the LNet activity map to enable direct comparison. RL deconvolution was performed in Python 3 using PyTorch.

For SID, we used the official SID implementation in MATLAB.^60^ The algorithm starts with background subtraction (2 iterations, detrending disabled) and computes a standard deviation image from the background-subtracted data. Initialization is then performed using non-negative matrix factorization with a rank of 15, an L1 norm regularization of the Gram matrix set to 1e-3, and 600 iterations. Cross-validation was enabled using 7 partitions. Each spatial component from the factorization is then reconstructed to obtain a volume, which is further segmented to extract neuron candidate positions. The reconstruction uses Projected Gradient Descent with Exact Line Search running for 3 iterations. L1 regularization was set to 0.1, and total variation regularization was applied with values 0.1, 0.1, and 4. Kernel optimization was disabled, and filtering was applied. The segmentation step was performed using a threshold of 0.1, with top and bottom cutoffs set to 5 and 49, respectively, and a neuron radius of 20 pixels. Cropping was not applied. All other parameters were set to their default values. Finally, the core of the SID algorithm consists of an iterative process alternating between spatial and temporal updates, refining both the temporal signals and spatial footprints, which constitute the detected neurons.

### Data analysis

We compared SNR across methods and as a function of depth below the slice surface. For the SNR calculation, the z-score of the time series was smoothed with a size 10 uniform filter. Noise (N) was calculated as the standard deviation of the difference between the smoothed and unfiltered traces. The signal (S) was the difference of the maximum and minimum values of the smoothed time series. SNR was calculated by taking the ratio and converted to a dB scale:

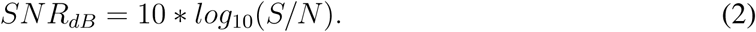

Time series were deconvolved using the oasis algorithm^86^ in the suite2P python module.^92^ Time-series correlations were measured using the Pearson correlation coefficient. Hierarchical agglomerative clustering on the deconvolved calcium time series sorted the neuronal responses into groups with similar activity. For each FOV the number of clusters was chosen based on the number of detected active neuron time series such that the average number of time series per cluster was ∼6.

Light-field processing and analyses were carried out in Python 3. Volumes and videos were visualized and rendered with napari.^93^

### Transfection and brain slice preparation

The following protocols were previously described in^68^ and summarized below.

This study was carried out in accordance with the recommendations of the UK Animals (Scientific Procedures) Act 1986 under Home Office Project and Personal Licenses (project license 70/9095). Mouse layer 2/3 cortical neurons were transfected by in-utero electroporation (IUE) at embryonic day (E)15.5 using soma-targeted jGCaMP8f (pAAV-CAG-RiboGCaMP8f^85, 94^) and tdTomato (pCAG-tdTomato, Addgene^95^). Time-pregnant female CD-1 mice (Charles River UK) were deeply anesthetized with 2% isoflurane. The uterine horns were exposed and intermittently rinsed with warm sterile phosphate-buffered saline (PBS). A plasmid mixture totaling 1–2 µg at a concentration of 1 µg/µl (6:1 ratio of jGCaMP8f:tdTomato), diluted in sterile PBS, was injected into one of each embryo’s lateral ventricles. Five electrical pulses (50 volts, 50 milliseconds each, at 1 Hz) were applied using 5-millimeter round plate electrodes (ECM 830 electroporator, Harvard Apparatus) with the anode positioned over the skull to target the cortex. Electroporated embryos were returned to the mother to continue development until birth.

Brain slices from electroporated mice were made at postnatal day (P)12–P30. 400-micron thick slices were cut in choline chloride solution containing (in mM): 110 choline-Cl, 25 NaHCO_3_, 20 glucose, 2.5 KCl, 1.25 NaH_2_PO_4_, 0.5 CaCl_2_ and 7 MgSO_4_. The slices were transferred to an artificial cerebrospinal fluid (ACSF) solution containing (in mM) 125 NaCl, 25 NaHCO_3_, 20 glucose, 2.5 KCl, 1.25 NaH_2_PO_4_, 2 MgSO_4_, 2 CaCl_2_, adjusted 300 to 310 mOsm/kg, pH 7.3 to 7.4 with HCl at 36C. Slices rested in ACSF for at least 20 minutes prior to transfer to a submersion chamber perfused with the same solution at room temperature throughout the imaging session. All solutions were oxygenated with 95% O_2_ / 5% CO_2_. Some slices were perfused with 50 µM 4-aminopyridine to evoke robust activity during imaging.

## Supplemental information

**Figure S1:** Calcium time series extracted from the same ROIs in volumes series reconstructed using LNet (yellow traces) and 8-iteration RL deconvolution (green traces).

**Figure S2:** Time series extracted from one light-field video via the SID algorithm showing the 6 dB inclusion criterion.

**Video S1**: Video of LNet (right), 8-iteration RL deconvolved (middle), and 1-iteration RL deconvolved (left) volumes, related to Figure 2.

## Data and Code availability

The raw light-field videos and LNet training weights are deposited at Zenodo [DOI: 10.5281/zen-odo.14900715].^96^ All original code for the end-to-end LNet Matrix and LNet ROI systems is publicly available at https://github.com/afoust/LNet jGCaMP8f pipelines.git.

## Materials availability

This study did not generate new unique reagents.

## Acknowledgements

This work was supported by the Wellcome Trust (311370/Z/24/Z), the United Kingdom (UK) Biotechnology and Biology Research Council (BB/R009007/1), the UK Engineering and Physical Sciences Research Council (EP/L016737/1, EP/W024020/1), and the Royal Academy of Engineering under the RAEng Research Fellowships scheme (RF1415/14/26). The authors thank Dr. Yu Liu for valuable assistance with transfection.

## Author Contributions

Conceptualization: CLH, KLYZ, HVJ, PLD, AJF; Data Curation: CLH; Formal Analysis: CLH, KLYZ, HVJ, AJF; Funding Acquisition: PLD, AJF; Investigation: CLH; Methodology: CLH, KLYZ, HVJ, PS, SJB, PLD, AJF; Project Administration: PLD, AJF; Software: CLH, KLYZ, HVJ, AJF; Supervision: PLD, AJF; Verification: CLH, KLYZ, HVJ, AJF; Writing – Original Draft: CLH, KLYZ, HVJ, AJF; Writing – Review & Editing: CLH, KLYZ, HVJ, PS, SJB, PLD, AJF.

