## Supplemental Figures for "Light-field deep learning enables high-throughput, scattering-mitigated calcium imaging"

**Supplemental Figure S1**

Calcium time series extracted from the same ROIs in volume series reconstructed using LNet (yellow traces) and 8-iteration RL deconvolution (green traces).

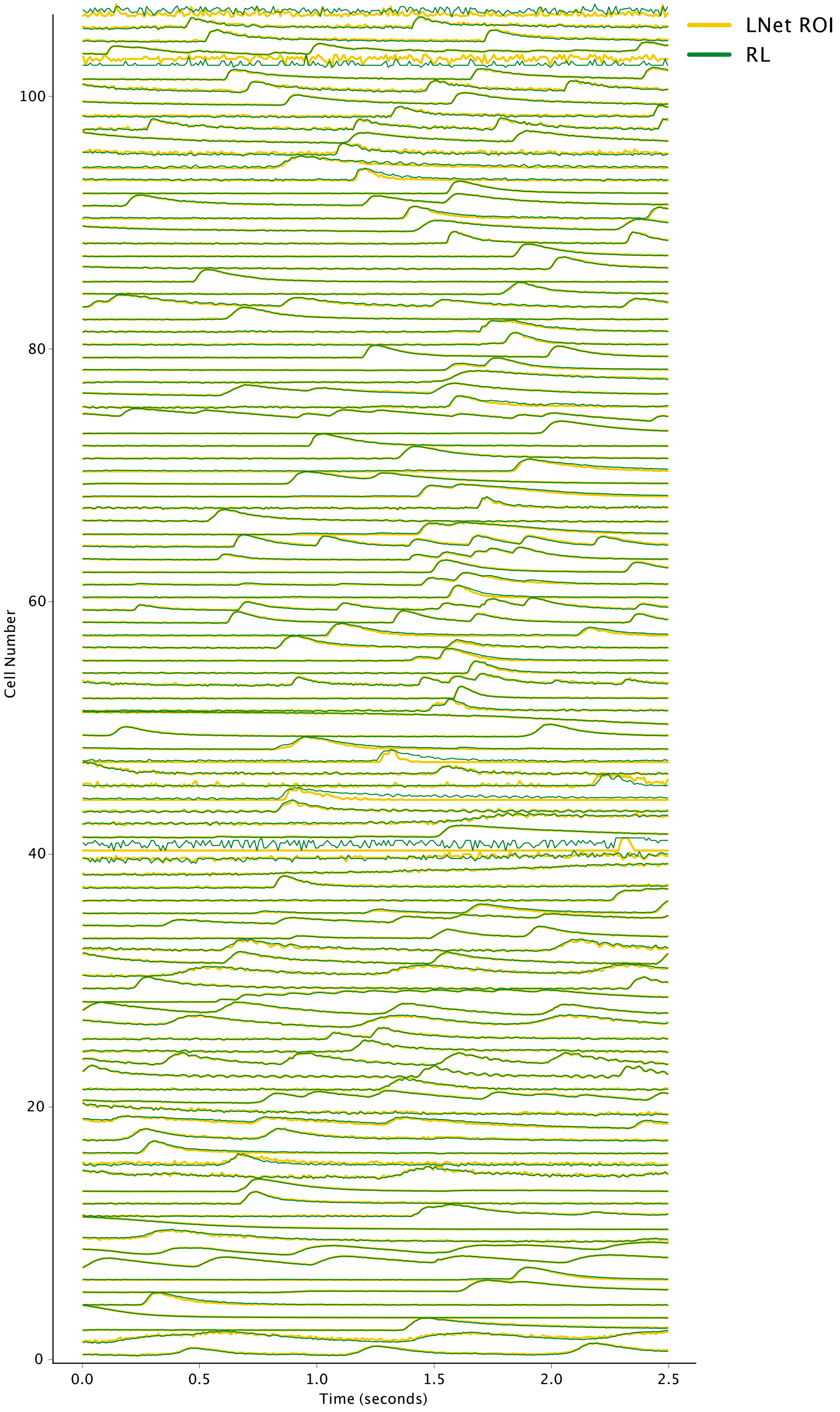

**Supplemental Figure S2,** choice of SNR threshold for time series inclusion:
Due to noise and ringing, only time series with SNR > 6 dB (red traces) were included in SNR and crosstalk analyses.

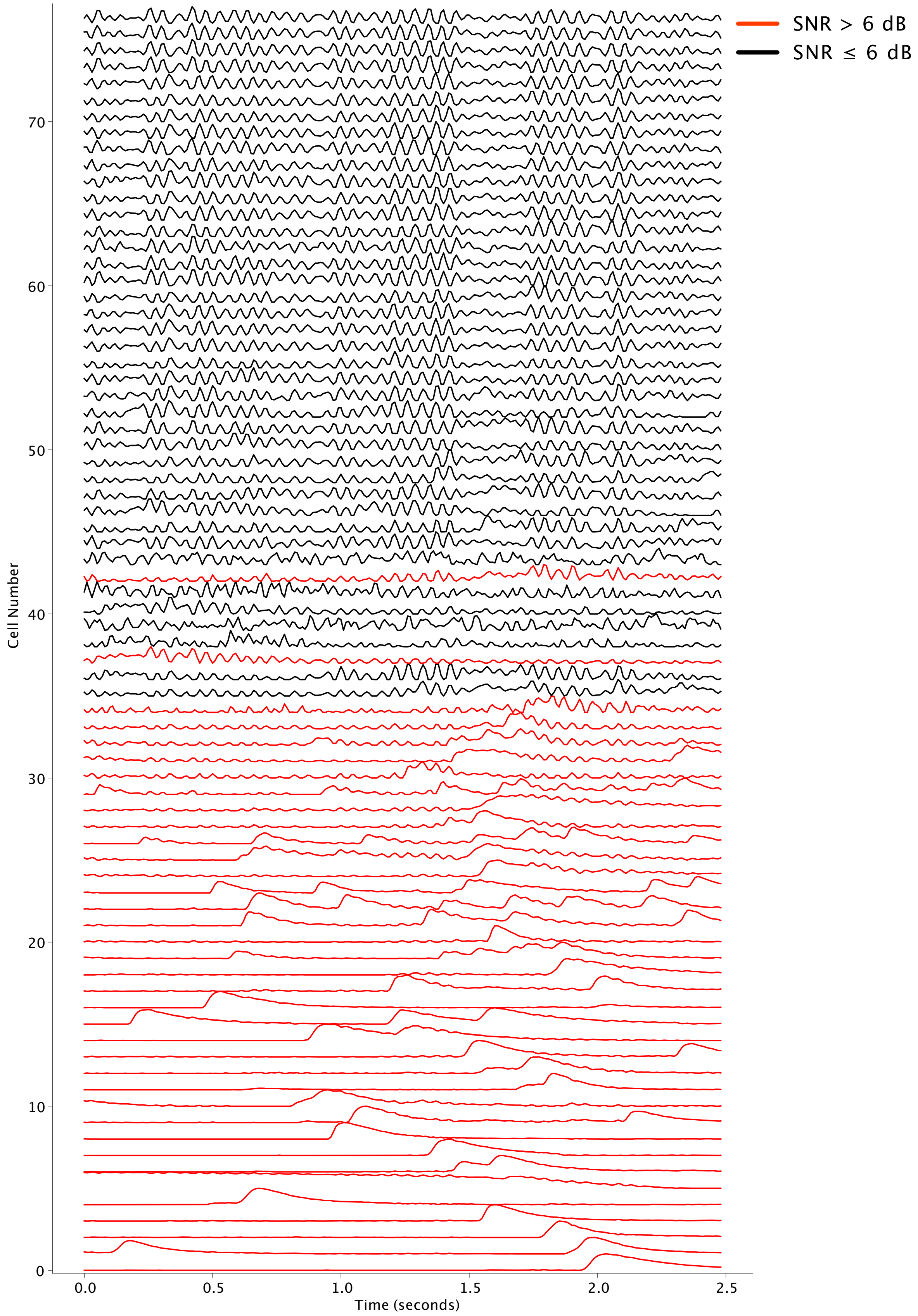

**Supplemental Video S1,** Video of LNet (right), 8-iteration RL deconvolved (middle), and 1-iteration RL deconvolved (left) volumes. The volumes are reconstructed from a 100 Hz light field video of ribo-jGCaMP8f expressing neurons in layer 2/3 neocortical slices. Each volume covers 533 x 500 x 104 microns displayed as x-y and x-z projections. The video plays at half real speed.
